# Fractionation-Free Protein Corona Quantification Through Synchrotron-Based Small-Angle X-ray Scattering

**DOI:** 10.64898/2025.12.19.695217

**Authors:** Juliana T. T. Carvalho, Caroline E. P. Silva, Lindomar J. C. Albuquerque, Antônio A. Malfatti-Gasperini, Liming Wang, Nathan P. Cowieson, Mateus B. Cardoso

**Affiliations:** Brazilian Synchrotron Light Laboratory (LNLS), Brazilian Center for Research in Energy and Materials (CNPEM), CEP 13083-100, Campinas, São Paulo, Brazil; Institute of Chemistry, University of Campinas (UNICAMP), CEP 13083-970, Campinas, São Paulo, Brazil; Laboratory for Interface Science and Biology, Institute of High Energy Physics, Chinese Academy of Sciences, Beijing 100049, P. R. China; Diamond Light Source Ltd, Harwell Science and Innovation Campus, Didcot, Oxfordshire OX11 0DE, United Kingdom

**Keywords:** Small-Angle X-ray Scattering, protein corona quantification, fractionation-free, synchrotron SAXS

## Abstract

When nanoparticles (NPs) enter biological environments, they are rapidly coated by biomolecules, forming the protein corona (PC) that defines their biological identity and dictates how NPs are recognized, distributed, and processed by living systems. Capturing the authentic features of the PC demands experimental conditions that preserve its native state, which are difficult to achieve once NPs are removed from their biological milieu. Despite significant progress, current PC quantification methods still rely on separating the NP-PC complex from its native environment, which compromises the corona’s integrity and prevents accurate evaluation of its physicochemical properties. Here, we introduce a fractionation-free approach based on synchrotron small-angle X-ray scattering (SAXS) to quantitatively determine the amount of protein adsorbed onto silica NPs under native conditions. By modeling the scattering contribution of free versus bound proteins, we directly extracted the adsorbed mass in both single-protein (BSA) and complex proteomic (human serum) systems. The resulting adsorption isotherms enabled the determination of thermodynamic parameters such as binding constants and cooperativity, distinguishing between monolayer and multilayer adsorption regimes. Together, these findings establish SAXS as a robust, non-invasive, and quantitative technique for probing the protein corona *in situ*, without perturbing the native equilibrium. This methodology paves the way for the development of new *in situ* analytical frameworks across diverse nanomaterial and proteomic systems, advancing SAXS toward quantitative characterization of the protein corona.

**Table of Contents Graphic:** 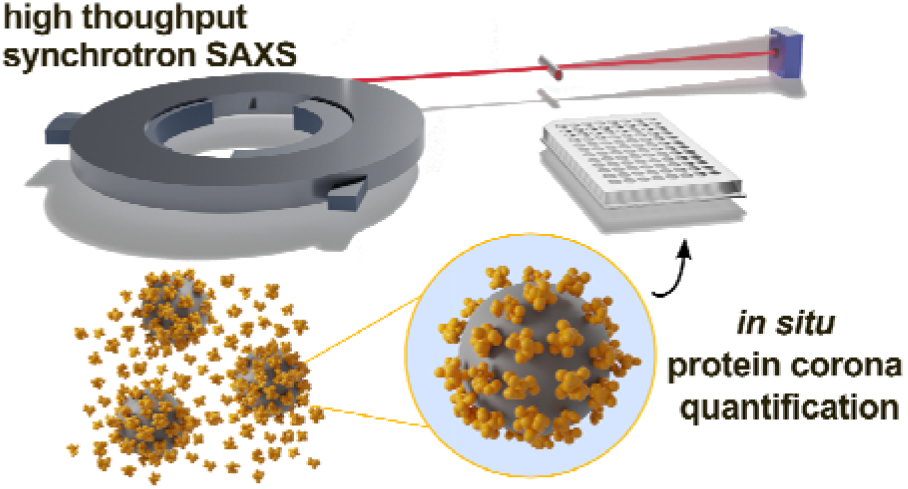

High-throughput synchrotron SAXS enables *in situ*, quantitative tracking of protein corona formation on nanoparticles.

## 1. Introduction

The notable increase in efforts to describe the formation of the protein corona (PC) is reflected in the growing number of studies that seek to manipulate, quantify, and understand this phenomenon under conditions closer to the natural biological scenario. ^1–3^ Reducing the translational gap between model studies and practical applications is crucial, as PC dictates possible outcomes for applying nanoparticles (NPs) in biological fluids.^4,5^ Consequently, experimental approaches that emulate real-world conditions are imperative to expand the current knowledge about PC and fast-track the effective use of NPs in various biomedical solutions, given the significant potential in this domain.

The PC is a well-established concept that ultimately describes an interface between the NPs surface and the surrounding media, formed mainly by proteins^6^ while other biocomponents (e.g., sugar, lipids, salts, and enzymes) can also be adsorbed. ^5,7,8^ PC is composed of hard and soft PC components, referring respectively to proteins that are strongly bound to the NP surface with high adsorption binding energy, and those that interact more loosely and can be displaced by components from the surrounding medium. These systems emerge from the almost immediate interaction between NPs and a fluid or biological entity, whose composition, thickness, and functional relevance may vary. ^9,10^ The PC is not a uniform entity, rather, it represents a complex interface shaped by both NPs properties and the biological environment and is influenced not only by the intrinsic properties of the NPs, such as size, shape, and surface chemistry, ^11–13^ but also by the characteristics of the adsorbed proteins. ^14^ These proteins may contribute differently to the composition of the PC depending on factors such as the presence of positive or negative domains, molecular size, abundance in the surrounding medium, and affinity for the particle surface. ^15,16^

Despite the significance of the PC, most characterization strategies rely on separating the NP–PC complex from its native medium. Classic fractionation*-*based quantification techniques include gel electrophoresis and mass spectrometry, which are trustworthy for obtaining corona’s proteomic composition and require hard corona separation from the native media. Common separation approaches, including centrifugation, filtration, magnetic separation, and chromatography, can modify the corona’s composition and structure, introducing artifacts that may not reflect the *in situ* state. ^17^ Recently applied techniques for PC probing, such as Size Exclusion Chromatography (SEC)^18^ and Asymmetric Flow Field-Flow Fractionation (AF4),^19,20^ promote a gentler separation of the corona but often demand extensive optimization and present low throughput. Labeling strategies (e.g., fluorescent tagging) can also modify protein conformation and charge, interfering with the thermodynamic and kinetic properties of adsorption. ^21^ A study using real-time single particle tracking was able to quantify the *in situ* soft corona of single, freely diffusing NPs under very low signal-to-background conditions, revealing that the total PC contained nearly twice as many proteins compared to values obtained from the extracted *ex situ* hard corona, in the exact same incubation conditions. ^22^ However, the requirement for dual labeling of both NPs and proteins ultimately constrains its applicability. Similarly, when the hard corona was captured *in situ* using an innovative “fishing” approach, its composition exhibited a strikingly low similarity (<30%) compared to that obtained by conventional centrifugation, ^23^ showing once more that isolation steps generate a corona that diverges almost entirely from its genuine *in situ* state.

Therefore, despite the plethora of available techniques for characterizing and quantifying the PC, those that rely on separating the NP–PC complex cannot be considered fully reliable. The separation process introduces unavoidable artifacts that distort both the total protein load and the spatial organization of adsorbed species. ^24^ For instance, centrifugation and magnetic separation can promote nanoparticle aggregation and entrapment of free proteins (i.e. proteins that did not constitute the PC), thereby influencing the detected corona. A comparative study demonstrated that the protein composition identified by these two techniques differed by more than 50%, highlighting that the choice of isolation method can substantially influence the reported corona profile. ^25^ Hence, characterization techniques that avoid fractionating the NP–PC complex from its native medium are essential, as they preserve the corona’s unique fingerprint and true physicochemical nature.^26^ Only *in situ* strategies can preserve key information that a faithful representation of the corona can obtain. Until now, there have been no reports in the literature demonstrating a direct and analytical method for quantifying the PC without the need for fractionation or separation procedures. In this scenario, Small-Angle X-ray Scattering (SAXS) has emerged as a powerful method for probing PC attributes and has already proven to be an adequate technique for investigating many aspects of the PC *in situ*, such as obtaining structural and thermodynamic features that are essential to describe a more realistic scenario of the PC. ^15,27,28^ SAXS offers high representativeness, rapid data collection, and the possibility to analyze NP-PC without prior modification. Importantly, all components of the native biological medium are preserved, which is particularly critical when studying interactions in complex fluids. Using synchrotron radiation as the light source in SAXS provides enhanced flux and signal-to-noise ratio, compared to lab-source SAXS, enabling a more efficient differentiation of the scattering contributions from NPs and their associated PC.

Herein, we present the first method capable of quantifying the PC without fractionating the NP–PC complex from its surrounding medium. We analyzed the SAXS scattering intensity, which is directly proportional to the mass concentration of the components in solution, allowing the extraction of the adsorbed protein mass in a fractionation-free and representative manner. We focused on evaluating protein adsorption onto silica NPs using two protein sources: bovine serum albumin (BSA) and human plasma (HS). Quantitative SAXS data provided measurable cooperativity and binding features for each system, which were best described by applying appropriate adsorption isotherm models (Langmuir, Freundlich, and Hill models). Additionally, the secondary structure of BSA upon interaction with NPs was further verified through synchrotron radiation circular dichroism (SRCD), ensuring that the overall integrity of the proteins remained intact throughout the process. While this study specifically examined BSA and HS on a single NP type, the quantification strategy is general and can be applied to a wide range of nanoparticle–protein systems. Overall, we present a fractionation-free quantification method capable of determining not only the total amount of adsorbed protein but also of providing a complete thermodynamic description that faithfully reflects the unperturbed state of PC, enabling the distinction between mono- and multilayer regimes in both single- and multicomponent systems.

## 2. Results and Discussion

High-Throughput Small-Angle X-ray Scattering (SAXS) experiments were done within the controlled environment of a synchrotron source at the B21 beamline of the Diamond Light Source - UK (**Figure 1a**).^29^ A robotic system was used to transfer each sample to the capillary cell, where measurements were taken to ensure precise and efficient handling. A critical consideration in this experimental setup was preserving sample integrity due to the high X-ray flux of the beamline and the biological nature of the specimen. Then, samples were prepared in 96-well plates, maintained at 37°C and the measurements were strategically carried out under carefully controlled flow conditions. This precautionary approach aimed to preserve the samples’ chemical and structural characteristics, thereby yielding accurate and reliable results that accurately represented their true properties. Each sample underwent a comprehensive measurement cycle consisting of 120 measurements, with each measurement duration lasting 0.25 seconds. This rigorous approach enabled a thorough examination of the samples’ attributes, contributing to the robustness and statistical significance of the overall findings. Initial measurements were carried out using thoroughly characterized pristine silica nanoparticles (SiO_2_) to evaluate the intrinsic intensity fluctuations within the measurement setup. SiO_2_ was selected for highly reproducible synthetic control over shape and size. ^30,31^ The primary objective of this phase was to assess the inherent variability in intensity readings for a singular sample. Then, a series of four successive measurement cycles was done on the same sample and the corresponding buffer, encompassing a process of capillary washing and drying between every measurement cycle. These repeated measurements revealed a maximum average deviation of approximately 0.3% from the obtained mean intensity value (**Figure 1b**), corresponding to a full amplitude variation in intensity of roughly 0.6% at low-*q*. These findings collectively underscore the stability and consistency of the measurement apparatus, demonstrating its capacity to maintain a high degree of precision even through sequential measurements.

**Figure 1.**
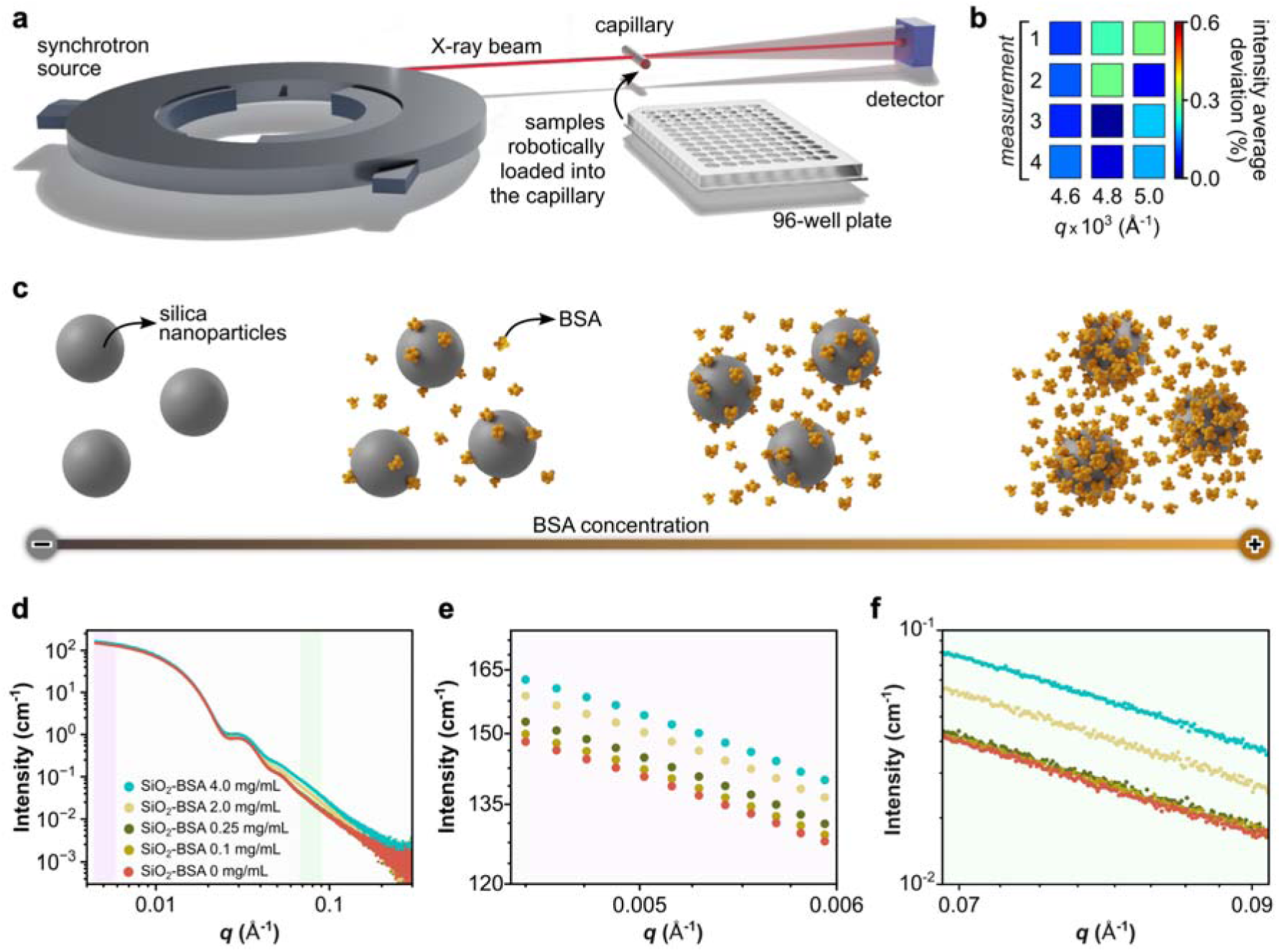
Nanoparticle-protein corona formation with bovine serum albumin (BSA) assessed by SAXS. (a) Schematic representation of SAXS experiments conducted in this study, emphasizing the synchrotron source and the 96-well plate. The samples were loaded into the capillary cell using robotics for the measurements. This schematic representation is not in scale. (b) Percentage intensity fluctuations at distinct *q* values observed over 4 consecutive measurement cycles to assess the precision achieved through sequential measurements. (c) Representation of SiO_2_ nanoparticles suspension before and after BSA mixing, indicating distinct samples containing varying protein coronas and excess free BSA. (d) SAXS profiles for pristine silica nanoparticles (SiO_2_-BSA 0 mg/mL) as well as for a few selected mixtures of SiO_2_ and BSA in PB media. The light purple and light green highlighted regions in panel (d) correspond to amplified areas in panels (e) and (f), respectively.

To explore the interaction of SiO_2_ nanoparticles within a biologically relevant context, the samples were incubated with BSA under controlled conditions. BSA was chosen as a model protein due to its close structural and physicochemical resemblance to human serum albumin (HSA) - the most abundant protein in human plasma and, sometimes, a constituent in the formation of nanoparticle-protein coronas. With comparable molecular weights, amino acid sequences, and tertiary structures,^32^ BSA serves as a robust and experimentally tractable biomimetic analogue for HSA, allowing for reproducible and mechanistically insightful investigations of nanoparticle-protein interactions.

During the incubation process, the SiO_2_ concentration was kept constant while the BSA amount was incrementally increased. (**Figure 1c**). The incubation was meticulously conducted to ensure a consistent reaction environment, and the samples were kept at 37°C before and during the SAXS measurements. Although the obtained scattering curves exhibit a remarkable similarity in their overall profiles (**Figure 1d**), a detailed analysis reveals substantial and meaningful distinctions, particularly in the regions highlighted in light purple and light green. Notably, at low-*q* values (light purple region - **Figure 1e**), pronounced differentiating features emerge that can be attributed to the PC effect and agree with the literature findings. This phenomenon is indicative of the BSA adsorption onto the SiO_2_ surface, resulting in alterations in their scattering behavior at low-*q*.^33^ Conversely, at high-*q* values (light green region - **Figure 1f**), the observed distinctions can be attributed primarily to the progressive increase of BSA excess (free proteins which were not adsorbed) within the system. It is essential to highlight that the shifts at low and high-*q* regions correlate well with the increase in BSA concentration. Therefore, the observed shifts in the scattering intensity behavior manifest with quantifiable differences between the SiO_2_-BSA 0 (control sample with no BSA) and 0.1 mg/mL, for example. Specifically, at low-*q* values, the shift amounts to a substantial ∼ 1.7 cm^-1^, while at higher-*q* values, the shift is more subtle at ∼ 0.002 cm^-1^. This discrepancy in shift magnitudes is particularly intriguing, as it directs attention to the potential influence of excess free BSA on the observed changes. However, the data robustly indicate that the pronounced intensity shift at low-*q* cannot be solely attributed to free BSA excess, since this magnitude at low-*q* surpasses what could be expected from the free BSA at equilibrium. Consequently, the PC formation must be considered the underlying driving force behind the observed significant shift at low-*q*, as previously reported in the literature.^33^ While we could argue that the magnitude of this intensity shift is larger but considerably close to what was presented in **Figure 1b** (intrinsic intensity oscillations related to the measurement), it becomes clear that with the escalation in protein concentration, the intensity shift amplifies and distinctly surpasses a 10% discrepancy. Importantly, the evolution of these scattering features follows a clear trend with increasing BSA concentration. Although minor oscillations in intensity could originate from intrinsic measurement fluctuations (**Figure 1b**), the systematic amplification of the low-*q* shift with increasing protein concentration (exceeding 10% beyond the baseline variation) strongly supports the conclusion that the observed signal originates from the PC formation, while the subtler high-*q* effects will be discussed later.^33^

To clarify the respective contributions of free BSA in solution and protein adsorbed onto the nanoparticle surface, we generated a series of theoretical scattering profiles that decompose the total SAXS signal into its fundamental structural components (**Figure 2**). This approach enables a direct comparison between the expected signatures of each contribution and the experimentally measured curves presented in **Figure 1**, thereby providing a mechanistic framework for interpreting the observed scattering evolution.

**Figure 2:**
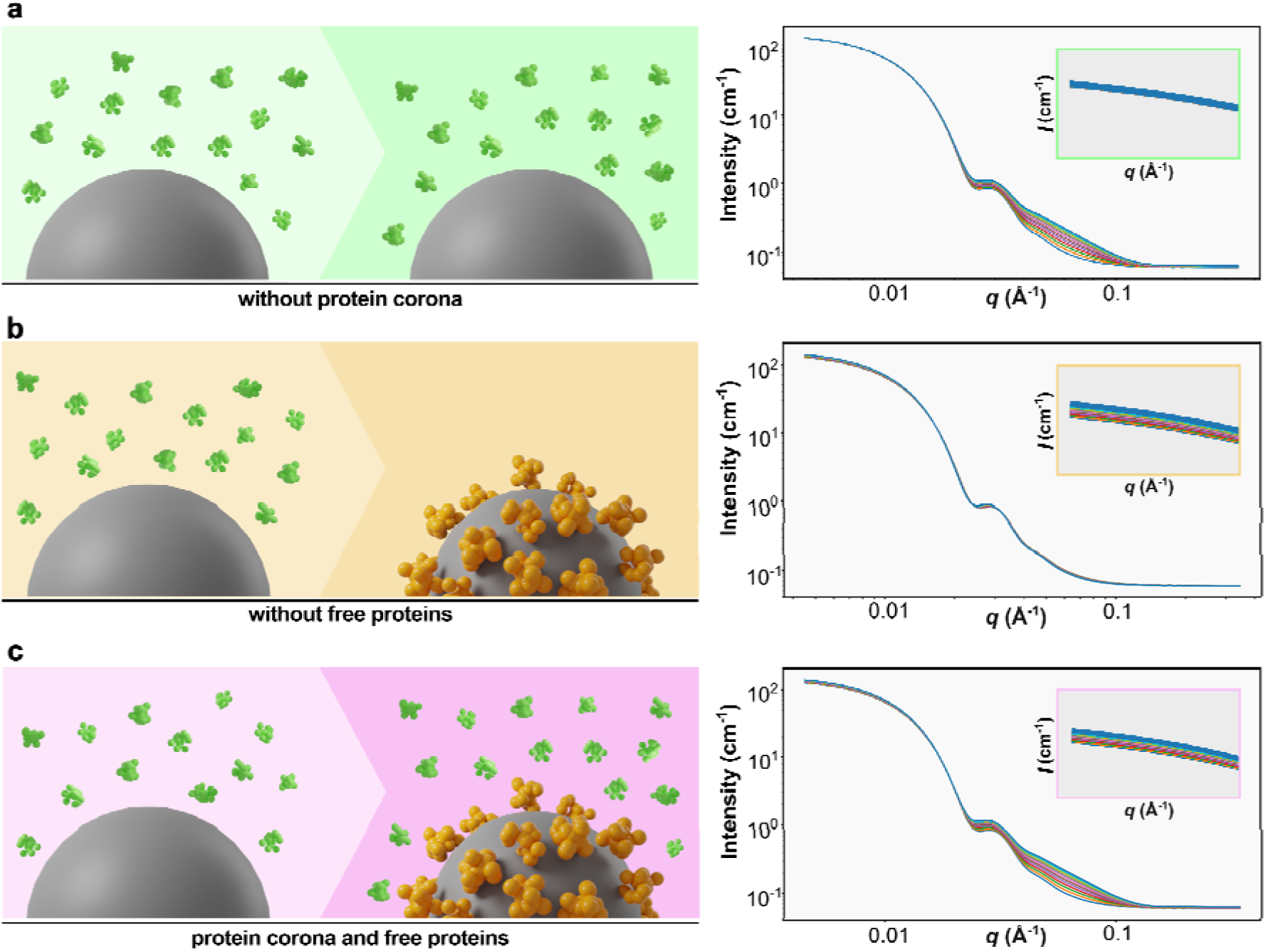
Modeled SAXS profiles illustrating the distinct contributions of free BSA and adsorbed protein corona. (a) Simulation including only free BSA, showing negligible changes at low-*q* and contributions confined to the high-*q* region. (b) Simulation including only the protein corona, yielding pronounced changes at low-*q* with minimal impact at high-*q*. (c) Combined simulation with free BSA and corona, reproducing the dual low-q and high-q features observed experimentally.

We began by characterizing the intrinsic scattering of the two principal constituents, silica nanoparticles and BSA, using independent form factor fits. The 400 Å SiO_2_ were well described by a polydisperse spherical model, yielding a mean radius of 174 Å and a radius polydispersity of ∼15%, consistent with their nominal dimensions. These fits employed the tabulated SLD of amorphous silica, ensuring accurate representation of contrast. In parallel, BSA was fitted as an ellipsoid using its theoretical SLD in buffer, resulting in polar and equatorial radii of 52 Å and 23 Å, respectively. This separate treatment captures the distinct length scales and scattering contrasts of the nanoparticles and the protein.

These simulations reveal distinct and diagnostically useful signatures across different *q*-regimes. When only free BSA is added to the model (**Figure 2a**), without any protein corona, the resulting profile shows little to no change in the low-*q* region, confirming that free protein contributes minimally at larger length scales. Conversely, when a corona is introduced in the absence of free protein (**Figure 2b**), the scattering is altered primarily in the low-*q* regime, with negligible influence on the high-*q* features, reflecting the smooth and relatively homogeneous nature of the adsorbed protein shell. When both effects are included simultaneously (**Figure 2c**), the composite curve reproduces the combined low-*q* and high-*q* trends observed experimentally, providing a direct mechanistic correspondence between the modeled and measured scattering behaviors.

Consequently, the simulations presented in **Figure 2** support a model in which the observed SAXS evolution arises from the combined influence of two concurrent processes: the accumulation of free BSA in solution and the formation of a protein corona around the silica nanoparticles. The model’s ability to capture both low-*q* and high-*q* behaviors provides strong evidence that corona formation alone cannot account for the experimental data, and that free protein contributes significantly to the overall scattering. This dual-component interpretation is consistent with the physicochemical expectations for nanoparticle–protein interactions and provides a mechanistic basis for the trends observed across the experimental dataset. Analytical models were generated using SasView. ^34^

**Figure 3a** presents the scattering profiles for three distinct samples: BSA (4 mg/mL), SiO_2_ (2 mg/mL) and SiO_2_-BSA mixture containing 4 mg/mL protein and 2 mg/mL silica particles. As expected, BSA curve diverges from the other SAXS profiles especially at low-*q*, while an overlay between the scattering patterns of pristine silica particles and those mixed with BSA is evident. However, the most pronounced deviation between these two silica-containing samples occurs at high-*q* values due to an excess of non-adsorbed BSA. **Figure 3b** highlights the disparities in the high-*q* region among the three scattering curves. Moreover, the introduction of a linear combination curve (black line) representing the summation of the scattering from pristine SiO_2_ and BSA surpasses the scattering intensity observed in the mixture of SiO_2_ with BSA. This observation highlights the non-linear nature of the interaction between SiO_2_ and BSA, indicating a scattering response that exceeds a simple additive combination. The difference, which is not accounted for in the model, relates to the presence of a protein corona on the surface of nanoparticles. We can subsequently fine-tune the weight of free BSA (**Figure 3c** - BSA excess) in the model to properly fit it with the experimental 10 curve, thereby determining the quantity of BSA that has become bound (**Figure 3c** - BSA corona).

**Figure 3.**
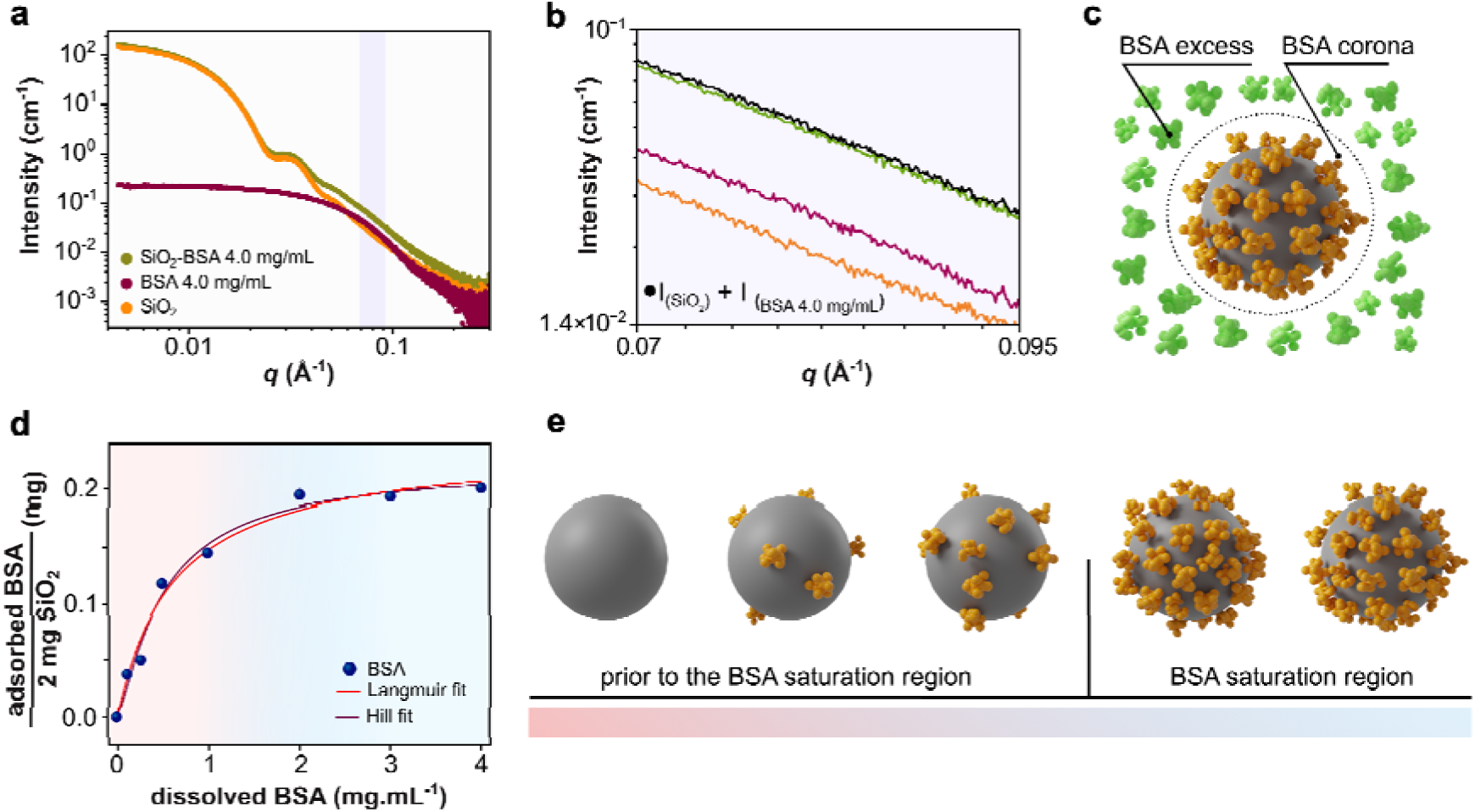
Quantification of BSA adsorption onto SiO_2_ nanoparticles. (a) SAXS profiles for pristine silica nanoparticles (SiO_2_-BSA 0%), BSA and a mixture of SiO_2_ and BSA 4.0 mg/mL in PB media. The purple highlighted region in panel (a) correspond to amplified areas in panel (b). (c) Schematic representation of a SiO_2_ nanoparticle surrounded by an adsorbed BSA corona and free excess proteins in solution. (d) Adsorption isotherm of BSA on SiO_2_, fitted with the Langmuir and Hill models, highlighting cooperative binding until reaching the saturation plateau. (e) Graphic representation of progressive BSA adsorption onto SiO_2_ nanoparticles, from the low-coverage regime to the saturation region, consistent with the experimental isotherm.

Using the experimental data, a binding curve was obtained by plotting the amount of adsorbed BSA (i.e., mg of BSA per 2 mg of SiO_2_) as a function of the free BSA concentration in solution (**Figure 3d**). The resulting profile was successfully fit by both Langmuir and Hill adsorption isotherm models. From these fits, we extracted a Hill cooperativity coefficient (*n_Hill_* = 1.23) and a Langmuir separation factor (*R_L_* = 0.13), both indicating a favorable SiO_2_/BSA interaction. ^35^ An in-depth discussion of the adsorption modeling is presented below. **Figure 3e** provides a graphical representation of the evolution of the BSA-corona, highlighting the region preceding saturation, rapidly reached at equilibrium BSA concentrations below 2 mg/mL, and the post-saturation regime observed at concentrations above this threshold.

To ensure that the structural features identified in the SAXS profiles truly reflect the formation of the protein corona rather than artifacts arising from possible protein degradation or conformational instability, we carried out a complementary structural investigation of the isolated corona. Given that SiO_2_ nanoparticles rapidly form a PC upon contact with BSA, it is essential to verify whether this adsorption event perturbs the native secondary structure of the protein, since the SAXS profiles obtained here primarily capture the scattering signal of proteins that retain their native conformation upon adsorption, rather than morphological alterations associated with unfolding or denaturation. To further validate this interpretation, Synchrotron Radiation Circular Dichroism (SRCD) spectroscopy provides a powerful means to simultaneously probe the formation of the corona and evaluate possible structural modifications of adsorbed proteins.

SRCD exploits the differential absorption of left- and right-circularly polarized light by optically active molecules, thereby offering sensitive insights into the absolute configuration and secondary structure of chiral biomolecules (**Figure 4a**).^36^ To ensure that the acquired CD signal originated solely from nanoparticle-bound proteins, the hard corona was isolated before measurement by removing unbound proteins through centrifugation and sequential washing steps. It is a mandatory procedure to eliminate spectral convolution from the free BSA, which could obscure the signal of the adsorbed layer (**Figure 4b**). The resulting CD spectra of the SiOL-BSA hard corona, obtained in PB, were compared with those of native BSA. The native BSA spectrum displays a characteristic maximum near 195 nm and two minima at approximately 209 and 222 nm,^37^ corresponding to its α-helical content (**Figure 4c**). The absence of significant spectral shifts in these features indicates that BSA maintains its secondary structure upon adsorption to the SiO_2_ surface under the studied conditions. This SRCD analysis confirmed that BSA preserves its structural integrity upon adsorption onto SiO_2_ nanoparticles, thereby validating that the observed scattering profiles primarily reflect nanoparticle–protein interactions rather than protein denaturation.

**Figure 4.**
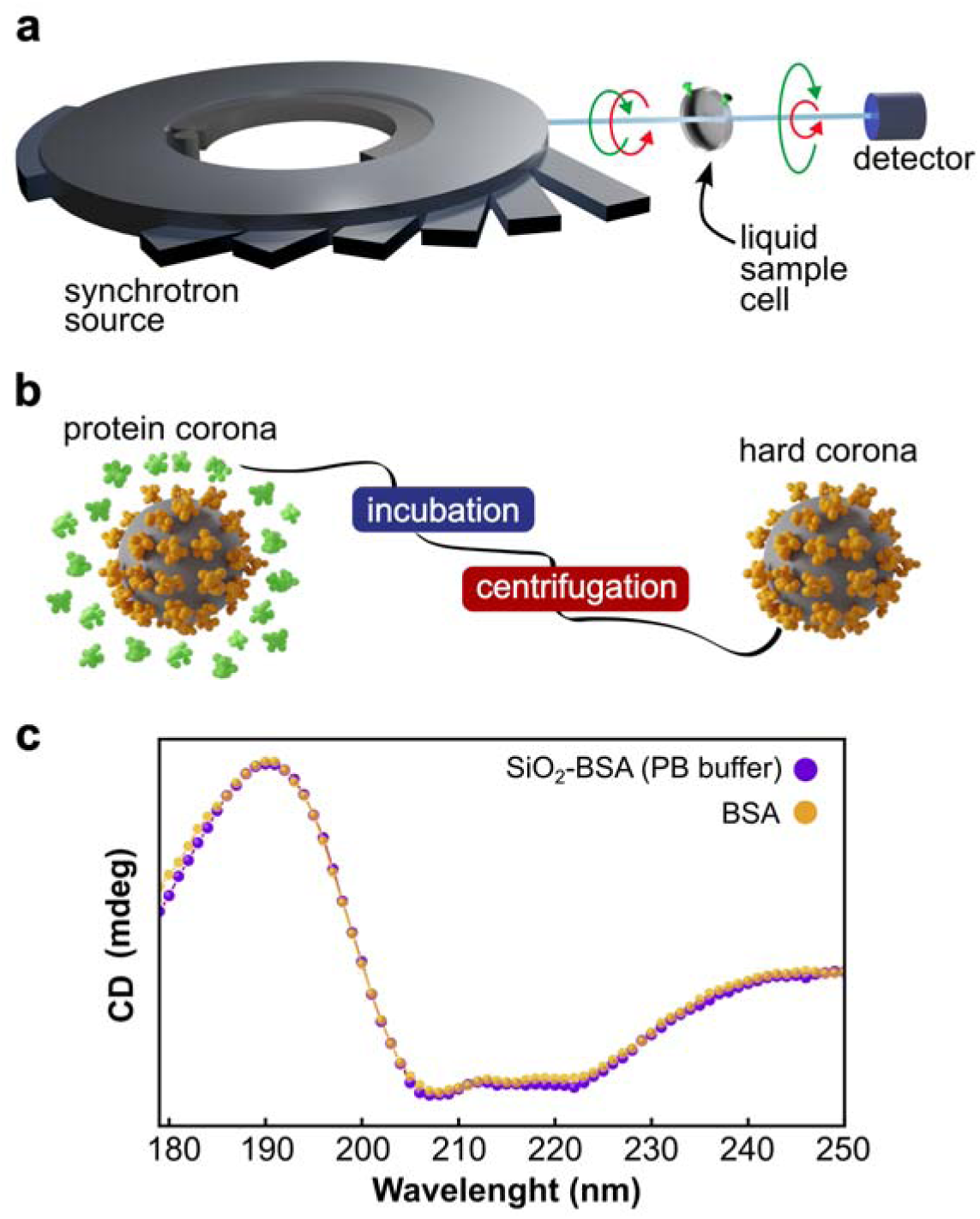
Synchrotron Radiation Circular Dichroism (SRCD) reveals that BSA retains its structural conformation upon interaction with SiO_2_. (a) Schematic representation of the SRCD spectroscopy setup, where circularly polarized UV light passes through the sample cell containing the SiO_2_-BSA solution. (b) Illustration of the separation process used to isolate the hard corona from free BSA in solution, which is critical to obtain SRCD spectra specific to the resulting corona (c) SRCD spectra of BSA in its native form (yellow), and after hard corona formation on SiO_2_ nanoparticles in PB (purple). The spectra reveal the preservation of the α-helical structure, as indicated by the characteristic minimum near 209 and 222 nm and a maximum around 195 nm.

SAXS can qualitatively assess the formation of a BSA-derived corona on SiO_2_ through an increase in scattering intensity at low-*q* regions, as previously discussed. This increase is attributed to the overall growth in particle size resulting from the adsorption of a protein layer, which directly correlates with the enhanced intensity at *q* values below 0.01 Å.^33^ To suppress corona formation, surface functionalization with antifouling moieties, such as zwitterionic (ZW) groups, emerges as a strategy to inhibit BSA adsorption on SiO_2_ surface. ZW groups have been extensively explored as hydrophilic coatings for NPs, forming robust hydration shells through electrostatic interactions with water molecules. ^38,39^

Here, SiO_2_ were functionalized with ZW groups, yielding a material with identical size and composition but featuring a ZW-functionalized surface (SiO_2_-ZW), which acts as a negative control for PC formation (**Figure 5a**). As a proof of concept, we exploited the antifouling properties of SiO_2_-ZW to provide a negative control for PC formation, thereby assessing how inhibited protein adsorption affects the SAXS profiles, more specifically at low-*q* (< 0.01 Å). SiO_2_-ZW particles were incubated with low, intermediate, and high concentrations of BSA (0.5, 2.0, and 5.0 mg/mL, respectively) under identical conditions of medium composition, incubation time, and temperature as previously used for pristine SiO_2_ (**Figure 5b**). The SAXS data for SiO_2_-ZW exhibit overall similar profiles to those of SiO_2_. Still, distinct features emerge particularly at high-*q* regions, where subtle differences in scattering intensity are evident and attributed to the progressive increase in free BSA within the system (**Figure 5c**). The effect of surface functionalization is apparent in the low-*q* region, where SiO_2_-ZW maintains stable scattering profiles with no intensity increase, confirming the suppression of BSA adsorption (**Figure S5**). This result contrasts with the scattering profiles of non-functionalized SiO_2_, where the presence of the PC leads to a pronounced increase in low-*q* intensity, as previously discussed and more clearly illustrated in **Figure 5d**.

**Figure 5.**
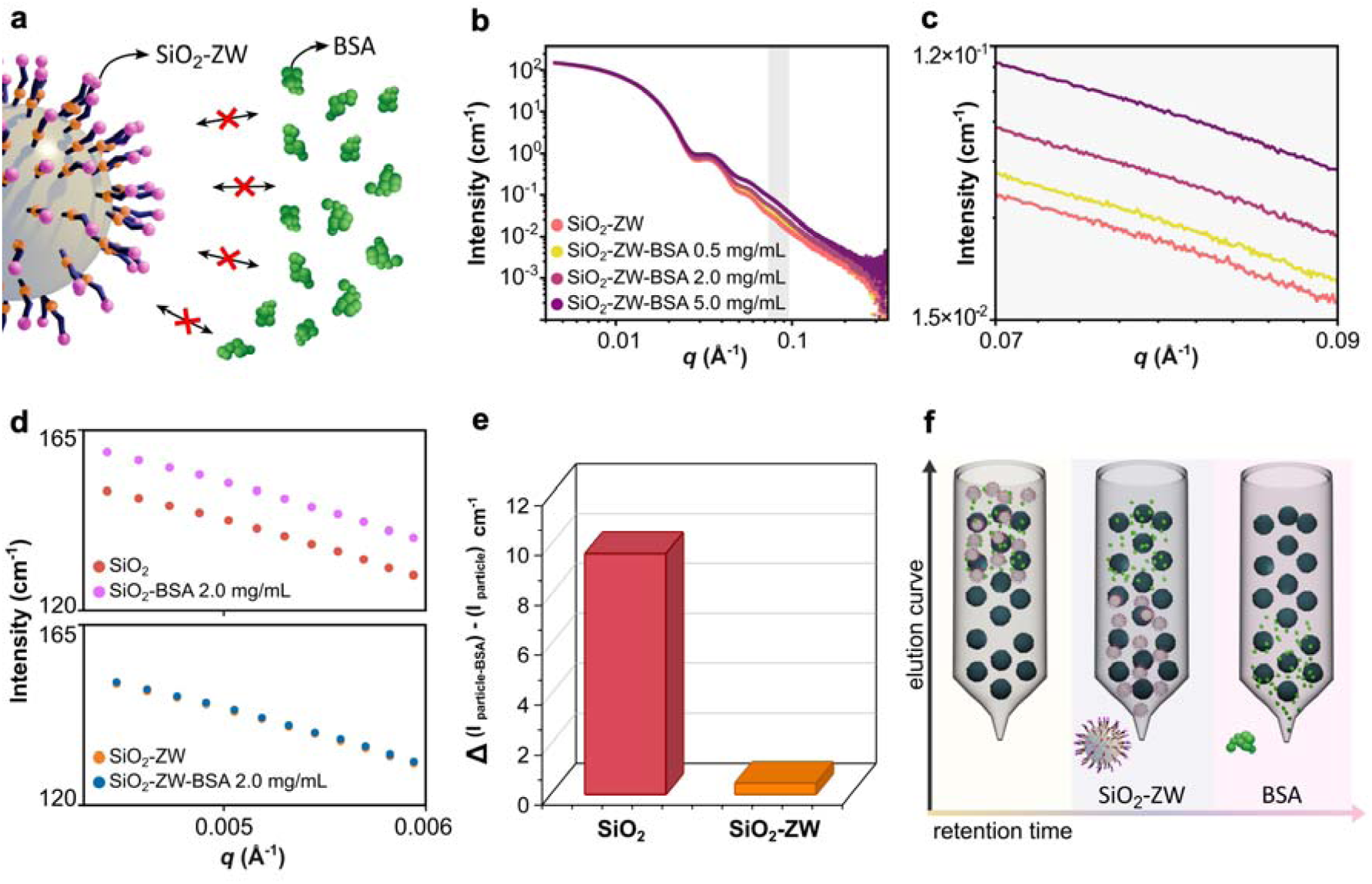
Negative control adsorption assay reveals distinct features in comparison with BSA-corona SAXS profile. (a) Schematic illustration of SiO_2_-ZW interacting with BSA, highlighting the suppression of protein adsorption due to the zwitterionic surface. (b) SAXS profiles for SiO_2_-ZW incubated with increasing BSA concentrations (0.5, 2.0, and 5.0 mg/mL), showing minimal changes in the low-*q* region. (c) Magnification of the high-*q* region from (b), evidencing the scattering contribution of the free BSA proportional to the increased concentration in solution. (d) Comparison of low-*q* intensity for SiO_2_ versus SiO_2_-ZW after incubation with 2.0 mg/mL BSA, confirming suppressed corona formation for SiO_2_-ZW. (e) Quantification of the intensity difference in the low-*q* region for SiO_2_ and SiO_2_-ZW, demonstrating the antifouling effect of the zwitterionic coating. (f) SEC-SAXS workflow used to separate nanoparticle–protein complexes from free BSA, enabling component-resolved scattering analysis.

These findings indicate that SiO_2_-ZW maintains excellent antifouling performance up to a BSA concentration of 2.0 mg/mL. At the highest concentration tested (5.0 mg/mL), a slight increase in scattering intensity around q ≈ 4 cm^-1^ is observed, suggesting that in this condition, the ZW functionalization is not sufficient to suppress BSA adsorption entirely. This likely indicates that not all the silanol groups in pristine SiO_2_ were modified by ZW, which can explain why, in a concentrated BSA regime, there is not a fully suppressed adsorption. Nonetheless, the distinct scattering behavior between pristine and functionalized NPs highlights that the analysis of low-*q* intensity is a reliable indicator of PC formation in the tested systems (**Figure 5e**). Moreover, as anticipated, it was not possible to quantify protein adsorption in this system as presented in **Figure 3d**, and consequently, an adsorption isotherm could not be obtained.

Furthermore, we performed an *in-situ* Size-Exclusion Chromatography–synchrotron SAXS (SEC–SAXS) experiment, which separates coexisting species by elution time and records their scattering profiles in real time.^29^ After incubating SiO_2_-ZW with BSA, the chromatographic run cleanly resolved the nanoparticle fraction from free protein (**Figure 5f**). Consistent with its antifouling character, SiO_2_-ZW eluted at a well-defined retention time distinct from that of BSA, and its SAXS profile lacked the excess low-*q* intensity characteristic of an adsorbed protein layer, behavior fully consistent with a “corona-free” state. In sharp contrast, bare (non-functionalized) SiO_2_ failed to elute under identical conditions since no nanoparticle SAXS peak was detected along the chromatogram. This strong retention is likely consistent with the formation of a BSA corona on the unmodified silica surface, which introduces additional interaction sites and promotes adhesive/bridging contacts with the chromatographic matrix, effectively immobilizing the particles. Taken together, the sharp elution of SiO_2_-ZW and the retention of bare SiO_2_ provide orthogonal evidence that the zwitterionic coating prevents corona formation, whereas the unmodified surface readily develops a BSA corona.

The fractionation-free quantification methodology demonstrated for SiO_2_-BSA mixtures was then extended by employing increasingly complex protein-based media, namely human serum (HS). Since the method proved suitable for a model system, e.g., a single-protein solution in PB buffer, it is natural to investigate its applicability in more complex scenarios, such as the one involving HS. HS is particularly relevant, as it represents one of the most important fluids for *in vitro* PC studies. ^40,41^ To assess these interactions, SiO_2_ nanoparticles were incubated with increasing percentages (v/v) of HS while maintaining a constant SiO_2_ mass across all conditions. The incubation procedure mirrored that used for SiO_2_-BSA, with meticulous control to ensure identical sample handling. Specifically, SiO_2_-HS was kept at 37 °C during both incubation and SAXS measurements. SAXS curves display the characteristic scattering profile of SiO_2_-HS particles, while detailed inspection of the low-*q* region reveals additional features attributable to protein adsorption (**Figure 6a and b**). For *q* < 0.01 Å^-1^, the scattering curve for HS mixture exhibits a steeper increase compared to that of pristine silica, consistent with subtle aggregation (more negative slope, typical of fractal-like or larger structures). Similar features have been reported in a similar scenario by Galdino et al. in studies of SiO_2_ interactions with complex proteomes, where the formation of dimers and trimers was observed.^33^ These results suggest a regime of subtle aggregation, rather than massive aggregates, is taking place. Importantly, the mass fraction of aggregates remains low and does not significantly impact the analysis of high-*q* data (**Figure 6c**), which are dominated by the contribution of free serum and remain valid for quantification purposes, even in the presence of minor NP aggregation.

**Figure 6.**
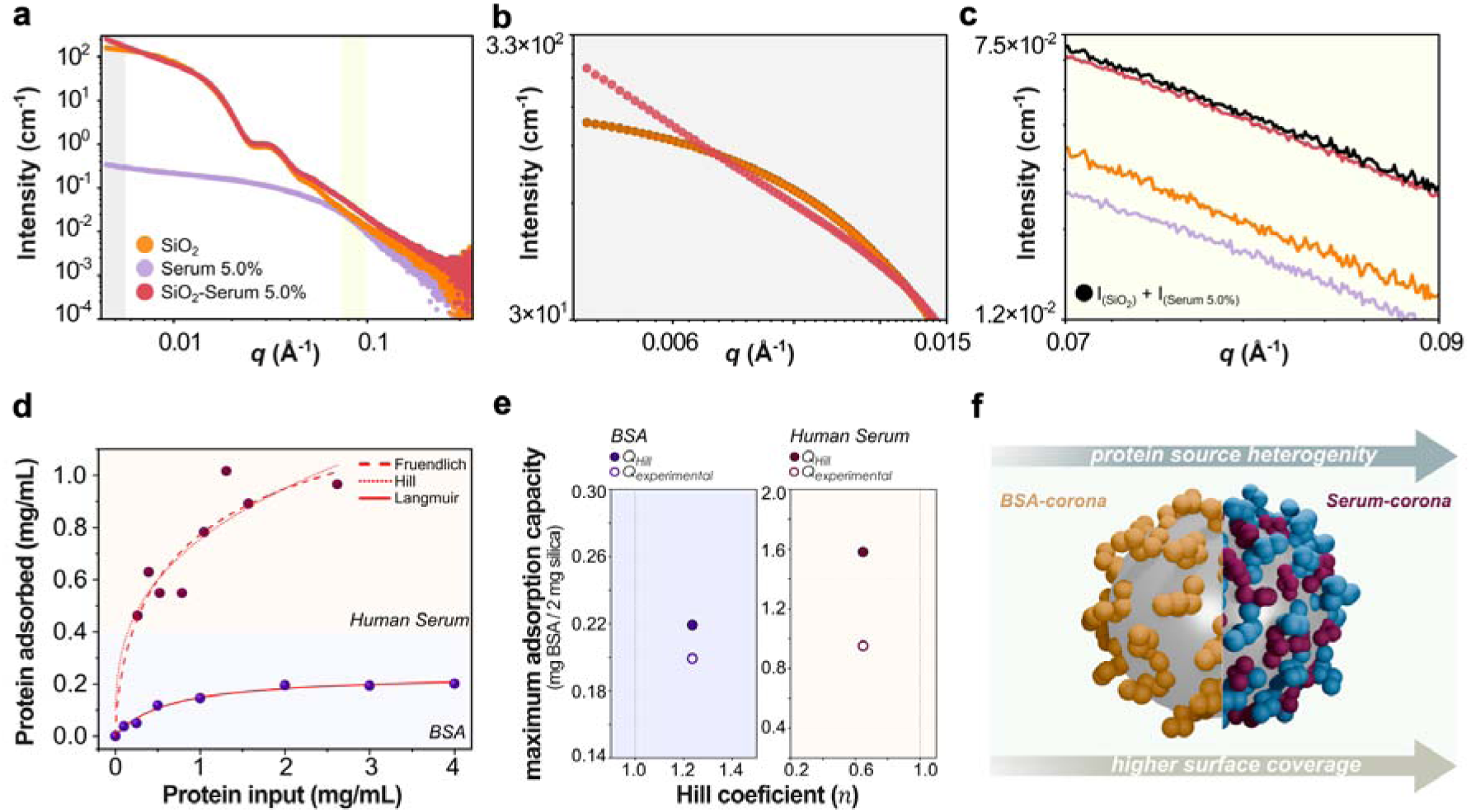
The protein corona assay for the SiO_2_-HS system reveals distinct features in both the SAXS scattering profiles and adsorption modeling results, indicating the formation of a more complex, multilayered corona structure. (a) Small-Angle X-ray Scattering (SAXS) profiles for silica nanoparticles (SiO_2_), Human Serum (HS, at 5.0%) and a mixture of SiO_2_-Serum 5% in PB media. The grey highlighted region in panel (a) corresponds to the amplified area in panel (b), relative to the low-*q* region. (c) Amplified area corresponding to the yellow highlighted region in (a), relative to the high-*q* region and demonstrating the sum of the scattering contribution from SiO_2_ and Serum 5.0% (black line). (d) Adsorption isotherms for BSA and HS experimental data, adjusted with Langmuir, Freundlich, or Hill models, evidence of a higher adsorption capacity for HS compared to BSA. (e) Plot of maximum adsorption capacity versus Hill coefficient for BSA and HS, illustrating the differences between experimental data and theoretical values predicted by the Hill model. The positioning of the data points in regions of positive (> 1) or negative (< 1) cooperativity highlights distinct adsorption behaviors for each protein regime. (e) Schematic illustration of distinct corona regimes: a single-protein system (BSA-corona), characterized by partial surface coverage of the SiO_2_ nanoparticle, versus a complex protein mixture (serum-corona), where the diversity of available proteins promotes enhanced and more complete surface coverage.

Adsorption equilibrium isotherm analysis offers a means to elucidate the relationship between the amount of protein adsorbed onto SiO_2_ surface and the concentration of unbound protein remaining in solution at equilibrium.^42^ More specifically, in this work, we aimed to saturate the NP surface under two distinct protein environments (BSA and HS). For each case, an increasing concentration series of the respective protein source was prepared, incubated and later measured at equilibrium under a defined contact time, temperature, and buffer conditions.

The adsorption extent was determined through absolute quantification, leveraging SAXS scattering intensities as a direct measure of the free protein concentration, with BSA employed as a representative model protein. The step-by-step quantification procedure, as described in the analysis of the SAXS curves, is outlined in the Supporting Information (**see Section S4**). The resulting isotherm profiles were then examined to assess the applicability of classical adsorption frameworks. In particular, the Langmuir, Freundlich, and Hill models were considered to distinguish the adsorption mechanisms in different protein sources.

Langmuir model implies that adsorption occurs in a monolayer until all surface-active sites are saturated, and can be described by the following non-linear equation:

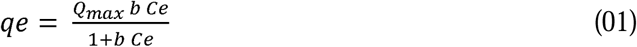

where *qe* is the mass of protein adsorbed per mL of suspension at equilibrium, with each mL containing 2 mg of silica nanoparticles; *Q_max_* represents the theoretical maximum adsorption capacity under these conditions (mg/mL); *b* is the Langmuir constant (mL/mg), which reflects the affinity between the protein and the nanoparticle surface; and *Ce* is the concentration of free protein in solution at equilibrium (mg/mL). The Langmuir equilibrium dissociation constant (*K_D_*) represents the equilibrium concentration of the adsorbate (mg/mL), in this case, proteins, that leads to half-saturation of the NPs surface, and can be obtained using the relation: *K_D_* = 1/*b*. Through *b*, it is also possible to obtain the separation factor (*R_L_*) which infers if the adsorption process is favorable (*R_L_ <* 1), irreversible (*R_L_*∼ 0), linear (*R_L_* = 1) or unfavorable (*R_L_* > 1). ^35^ This parameter is obtained by the following relation:

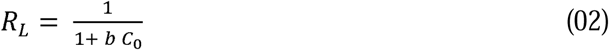

where *C*_0_ is the highest initial concentration of protein.

The Freundlich isotherm model assumes a heterogeneous surface with high and low affinity regions, but with a progressive reduction of adsorption affinity due to lateral repulsion between the adsorbed molecules. The model equation is represented as follows:

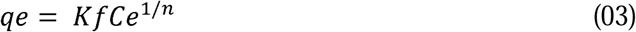

where *Kf* is the Freundlich constant, related to the adsorption capacity of the system. Higher values indicate a greater amount adsorbed for a given concentration of free protein and *n* is the exponent of the Freundlich model, where 0 < *n* < 1 is favorable adsorption, *n* = 1 indicates linear isotherm, and *n* > 1indicates cooperative adsorption.

Lastly, the Hill model describes adsorption as a cooperative process, in which the binding of an adsorbate at one site on the adsorbent affects the affinity of other available sites on the same surface. The Hill coefficient (*n_Hill_*) represents the degree of cooperativity: *n* > 1 indicates cooperative binding, *n* < 1 indicates negative cooperativity. ^3,33^ The model equation is expressed as follows:

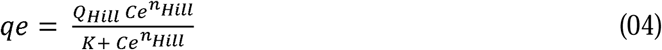

where *Q_Hill_* is the theoretical maximum adsorption capacity (mg/mL), *K* is the Hill constant, and *n_Hill_* is the dimensionless Hill coefficient. Additionally, the effective dissociation constant (*K*′*_D_*) corresponds to the midpoint concentration of the adsorption isotherm and was obtained from the Hill parameter *K* using the relation *K*′*_D_* = *K*^l/*n_Hill_*^.^41^ The judgment of the best fit was based on the values obtained by the nonlinear Chi-Square Test (χ^2^) and the coefficient of determination (*R*^2^). ^43^ All of the obtained adsorption parameters for BSA and HS are presented in **Table S2** and the obtained isotherms are shown in **Figure 5d**.

The adsorption isotherms obtained for BSA on SiO_2_ were fitted using Langmuir, Freundlich, and Hill models. The Langmuir model provided an excellent fit to the experimental data (*R*^2^ = 0.9837, χ^2^ = 1.2 × 10^-4^), yielding a maximum theoretical adsorption capacity (Q_max_) of 0.239 mg/mL and an affinity constant (*b*) of 1.61 ± 0.29 mL/mg (**Figure 5d, lower panel**). Similarly, the Hill model also described the data with high accuracy (*R*^2^ = 0.986), returning a maximum coverage (Q_Hill_) of 0.22 mg/mL, in close agreement with the Langmuir estimate. This convergence between the two models indicates that adsorption occurs predominantly as a compact layer, consistent with an effective monolayer regime. In our model system, the BSA adsorption isotherm (blue region) shows a steep increase in surface-bound BSA for equilibrium concentrations (*Ce*) below 2 mg/mL, indicating strong attractive interactions between BSA and SiO_2_. Above this threshold (*Ce* > 2 mg/mL), the isotherm reaches a plateau, corresponding to the maximum surface coverage that can be achieved under our experimental conditions. In this saturation regime, increasing the bulk protein concentration from 2 to 4 mg/mL did not increase the amount of adsorbed BSA, which remained constant at ∼ 0.20 mg/mL. This experimental value is in close agreement with the *Q_max_* values predicted by Langmuir (0.23 mg/mL) and Hill (0.22 mg/mL) model fits, demonstrating the robustness of the isotherm modeling. Additionally, the equilibrium dissociation constant (*K_D_*) for Langmuir and the effective *K_D_* for Hill were 0.61 and 0.5 mg/mL, respectively.

It is well established that BSA corona formation does not fully cover the total available nanoparticle surface area. Even under saturation conditions (protein excess in solution), adsorption is typically limited to a compact layer with a density close to, but not exceeding, that of a complete monolayer. This limitation arises from protein–protein interactions, which influence the packing arrangement and prevent perfect surface coverage. For BSA, with an isoelectric point of approximately 4.7, not significantly far from the pH used in our experiments (∼7.0), the net negative charge is moderate, allowing electrostatic attraction through localized positive patches to overcome the overall repulsion and promote adsorption. However, the interplay of these electrostatic effects with steric constraints and intermolecular interactions ultimately restricts surface coverage to less than 100%, which can explain why, in these experimental conditions, the full coverage of the SiO_2_ surface is not achieved. Another evidence that supports the hypothesis that a monolayer is occurring for the SiO_2_-BSA adsorption process is the lack of significant improvement in the Freundlich fit, which showed lower agreement with the data (*R*^2^ = 0.9081, χ^2^ = 5.23 × 10^-4^) in comparison with the previous models.

In contrast, HS exhibited markedly higher overall adsorption and affinity toward the silica surface, reflecting the complex mixture of proteins present in this biological fluid. The Freundlich model provided a reasonable fit (*n* = 3.01, *R*^2^ = 0.7649, χ^2^ = 0.012), but the Hill model more accurately captured the adsorption behavior (*R*^2^ = 0.9106), yielding a maximum adsorption capacity *Q_Hill_* = 1.6 ± 1.8 mg/mL and a cooperativity coefficient *n* = 0.64 ± 0.66. The sub-unity Hill coefficient indicates negative cooperativity and competitive binding, consistent with the idea that early adsorption events hinder subsequent access of other proteins to available surface sites. The absence of a clear saturation plateau suggests that adsorption continues beyond a single monolayer, which explains the large uncertainty associated with the estimated maximum capacity.

The superior Hill fit implies that heterogeneous, multi-protein adsorption processes dominate in HS, rather than uniform surface binding as described by the Freundlich model. This behavior likely arises from the coexistence of proteins with distinct affinities and sizes, such as immunoglobulins, fibrinogen, and apolipoproteins, which compete for surface sites and can also engage in protein–protein interactions, promoting secondary layer formation. Similar multilayer adsorption regimes have been reported in serum-nanoparticle systems, supporting this interpretation.^44^ Importantly, this scenario suggests that not all proteins in the corona are in direct contact with the silica surface; rather, a hierarchical architecture emerges, where early bound species modulate the subsequent adsorption of others, defining the compositional and structural heterogeneity of the corona.

For instance, a comparative analysis of the Freundlich constants (*Kf*), which represents the relative adsorption capacity while considering the same available SiO_2_ surface area for interaction, reveals that SiO_2_ exhibits a lower affinity for BSA (*Kf* = 0.13) compared to human serum (*Kf* = 0.60). These results are consistent with the experimentally determined adsorption capacities (**Figure 5e**), where HS (0.96 mg/mL) displays a value approximately five times higher than that of BSA (0.20 mg/mL), further corroborating the correlation indicated by the *Kf* parameters. Also, the Hill coefficient (*n*) differs between the single-protein (BSA) and multiprotein (human serum) systems, indicating positive cooperativity in the latter and negative cooperativity in the former.

Taken together, these results indicate that HS gives rise to a corona that continues to grow, making it non-trivial to obtain a saturation plateau under the tested conditions. In particular, the obtention of the *K_D_* parameter is limited in HS scenarios, mainly due to the highly heterogeneous nature of these protein sources. Given that serum concentrations up to 5% were tested, it is possible that the protein corona was still incomplete, leaving the NP surface partially exposed. Such partial coverage could create a heterogeneous interface that combines hydrophilic domains from silica with hydrophobic patches from adsorbed proteins, thereby reducing colloidal stability. Importantly, in HS, it is often difficult to evaluate whether the adsorbed proteins undergo conformational denaturation, thereby exposing hydrophobic domains that may promote secondary protein-protein interactions and subsequent NP aggregation. In multiprotein systems, this complexity is further amplified, as techniques such as SRCD still face limitations in resolving overlapping spectral contributions.

At higher serum concentrations (above 5%, not assessed here and out of the scope of this work), a more continuous protein layer would likely form, potentially restoring colloidal stability through steric and electrostatic repulsion. Further work is needed to validate this assumption. Despite the occurrence of some aggregation, our methodology demonstrated sufficient robustness to allow quantitative analysis and extraction of thermodynamic parameters. This robustness contrasts with *in situ* techniques such as Fluorescence Correlation Spectroscopy (FCS), which are unsuitable for aggregating systems since even a small number of aggregates can dominate the autocorrelation function and obscure the signal from individual NPs.^45^

Hence, the adsorption parameters obtained here reflect a conditional adsorption regime, meaning they are intrinsically dependent on factors such as contact time, temperature, and medium pH. Here, we do not intend to maximize the adsorption capacity of SiO_2_, but rather to demonstrate that such parameters and an overall system description can be extracted directly from equilibrium SAXS data under native *in situ* conditions. By enabling quantification of adsorption without corona separation, this approach allows the discrimination between monolayer and multilayer regimes (**Figure 6f**), reinforcing SAXS as a direct and *in situ* characterization tool.

## Conclusion

Here, we present a fractionation-free, high-throughput SAXS methodology that enables quantitative analysis of nanoparticle–protein interactions in solution. This approach extracts binding affinities and cooperative effects while circumventing the well-known artifacts introduced by centrifugation or separation protocols. SAXS proved capable of robustly characterizing corona formation in both simple (BSA) and complex (human serum) environments, allowing us to derive thermodynamic parameters through isotherm modeling. The BSA-corona followed the Langmuir isotherm, confirming the predominance of a monolayer regime, while the human serum-corona was best described by the Hill model, revealing positive cooperativity and the formation of multilayers, consistent with the absence of a saturation plateau. The theoretical adsorption capacities (0.23 and 0.22 mg/mL for Langmuir and Hill, respectively) for BSA were in excellent agreement with the experimental value (0.20 mg/mL). At the same time, for HS, the Hill model predicted a higher apparent capacity (1.59 mg/mL) compared to the experimental observation (0.96 mg/mL), reflecting the challenge of achieving full saturation in multicomponent systems where layer stacking occurs. Within this scenario, SAXS stand out as a suitable option for a non-invasive quantification method, marking a significant step toward more reliable and physiologically relevant nanoparticle characterization.

## Acknowledgements

M.B.C. and L.W. acknowledge the support of the China-Brazil Joint Laboratory for Synchrotron Science and Technology. Authors are acknowledged to Fundação de Amparo a Pesquisa do Estado de São Paulo (FAPESP) (grants n° 2021/12071-6, 2023/13208-0, 2023/02144-1, and 2024/00989-7). The authors would like to thank the CEDRO beamline staff of the Brazilian Synchrotron Light Laboratory (LNLS) (proposal #20241402) for their assistance during the analyses. This work made use of access to Diamond Light Source and was supported by iNEXT-Discovery, funded by the European Union’s Horizon 2020 programme (PID: 26126). We also acknowledge Diamond Light Source for access and support at beamline B21 (proposal SM35502).

