## Supplementary Information for "Fractionation-Free Protein Corona Quantification Through Synchrotron-Based Small-Angle X-ray Scattering"

### **SECTION S1: Synthesis and Functionalization of SiO_2_**

#### *Materials*

Tetraethyl orthosilicate (TEOS, 98%), ammonium hydroxide (28-30%), sodium phosphate dibasic (Na_2_HPO_4_) and bovine serum albumin (BSA), anhydrous ethyl acetate, N,N-dimethyl-3-aminopropyltrimethoxysilane and 1,3-propanesultone.were purchased from Sigma-Aldrich (Brazil). Human serum (HS) from human male AB plasma (USA origin, sterile-filtered) was purchased from Sigma-Aldrich (UK). Ethanol (≥ 99.8%) was purchased from Merck. All the solutions and suspensions were prepared with ultrapure water obtained from a water purification system (Purelab from ELGA - resistivity of 18.2 MΩ∙cm). All reagents were used as received.

#### *Synthesis of Silica Nanoparticles (SiO_2_)*

Pristine silica nanoparticles (SiO_2_) were synthesized using a modified Stöber method, already well established in our group. ^1,2^ In a precisely controlled environment, 120 mL of ethanol and 4 mL of NH_4_OH were added into a flat-bottom flask and maintained under constant magnetic stirring at room temperature. After 30 minutes, 2.5 mL of TEOS was added into the solution and allowed to react while continuously stirring. Subsequently, after a monitored 3-hour period, a second aliquot of 2.5 mL of TEOS was added dropwise, and the system was subjected to extended overnight stirring under stringent conditions. The resulting suspension was subjected to a purification process involving dialysis, employing a cellulose dialysis membrane with a selected cutoff of 14 kDa. This synthetic route ensured the production of highly monodisperse SiO_2_ for the subsequent experiments. Initially, a previously prepared 4:1 ethanol-water mixture (v/v) was employed as the dialysis solution, and a rigorous protocol was followed. Subsequently, a transition was made to employing exclusively deionized water as the dialysis medium. The dialysis solution was replaced at 2-hour intervals until the pH level of the suspension reached a monitored 6.0, at which exact point the process was halted. The SiO_2_ final concentration was determined through gravimetry, and this procedure was conducted in triplicate (n = 3) to ensure accuracy. The resultant SiO_2_ aqueous suspension was then stored under temperatures between 4-8 °C, maintaining its integrity and quality for future utilization.

#### *Functionalization of SiO_2_ with Sulfobetaine Silane (SiO_2_-ZW)*

The pristine SiO_2_ nanoparticles were functionalized with sulfobetaine silane (SBS) zwitterionic moieties, following an adapted protocol from Litt et al.^3^  Firstly, the ligand was synthesized, by mixing anhydrous ethyl acetate (30 mL), N,N-dimethyl-3-aminopropyltrimethoxysilane (5 g, 2.173 mL) and, after 5 minutes, 1,3-propanesultone (3 g, 0.912 mL) under flux of N_2_ in a two-neck round bottom flask and the reaction was carried under constant stirring at 45 °C. After 2 hours, more ethyl acetate (10 mL) was added and the N_2_ flow was stopped. The heating ceased 3 hours later, and the reaction was maintained for 15 hours. The reaction mixture was redispersed in acetone and centrifuged at 5000 rpm for 5 min at 20 °C. This process was repeated 4 times. Finally, the product (SBS) was vacuum dried and stored in a desiccator. In the second step, SBS (100 mg dissolved in 10 mL of water) was added to SiO_2_ suspension (50 mL) under stirring, in a round bottom flask. After 15 min, NH_4_OH (2 mL) was added into the flask that was heated at 80 °C for 4 h. Then, the modified nanoparticles with SBS were washed three times with Milli-Q water followed by centrifugation at 10.000 rpm for 15 min at 10 °C. Finally, the functionalized nanoparticles were purified by washing and dialysis (cellulose dialysis membrane with a cutoff of 14 kDa) against ultrapure water. The suspension final concentration was determined by gravimetry, measured in triplicate (n=3). The SiO_2_-ZW aqueous suspension was stored at 4-8 °C for further use.

### **SECTION S2: Nanoparticle Characterization**

#### *Electron Microscopy*

The size and morphology of SiO_2_ nanoparticles were evaluated using scanning transmission electron microscopy (STEM) in secondary electrons and bright field modes. Images were acquired with a Thermo Fisher Scientific Scios 2 DualBeam operating at 30 kV. A 5 μL aliquot of the sample, previously dispersed in an ultrasonic bath for 10 minutes, was deposited onto a carbon-coated grid (400 mesh, TED PELLA®), and the solvent was allowed to evaporate for 10 minutes. Excess liquid was removed using filter paper. The size distribution of the NPs was determined by calculating the average diameter from a count of more than 250 nanoparticles and selected images are shown in Figure S1.

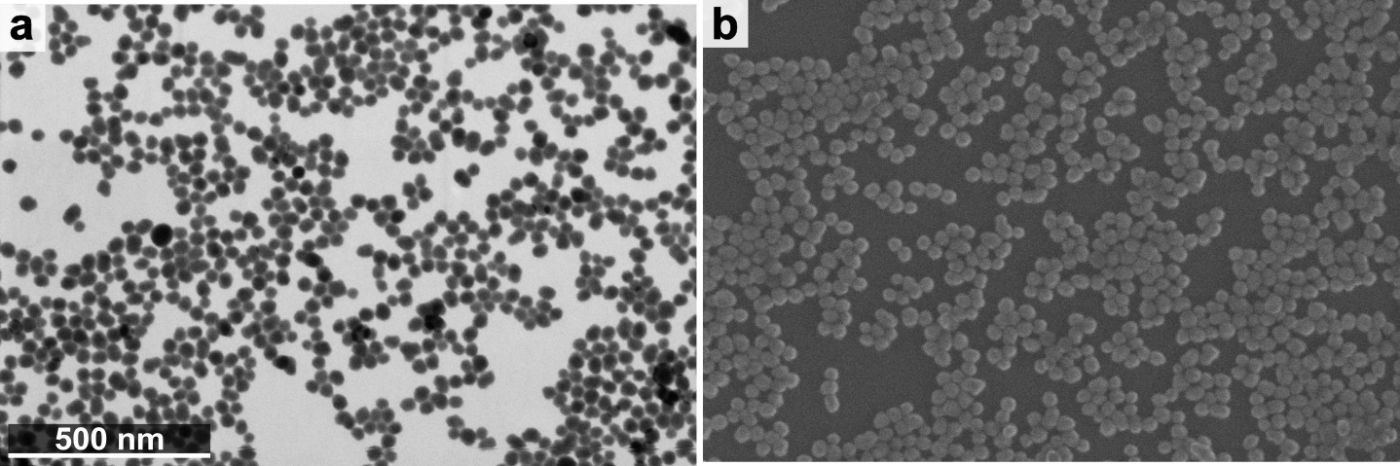

**Figure S1.** Morphological characterization of SiO_2_. STEM (a) and SEM (b) micrographs of pristine SiO_2_, with scale bars of 500 nm.

#### *Dynamic Light Scattering (DLS)*

The hydrodynamic diameter (D_H_) of pristine and functionalized SiO_2_ were determined using Dynamic Light Scattering (DLS) with a Zetasizer Malvern Nano ZS instrument (Malvern Instruments Ltd., UK). The instrument employed a red laser (632.8 nm) in backscatter mode (detection angle = 173°). A 2 mg/mL suspension of both particles were prepared in PB and ultrapure water for D_H_ and zeta potential (ζ) analyses, respectively. The measurements were performed in triplicate, where each measurement consisted of 10 runs of 10s at 25 °C with thermal stabilization of 120s. The results obtained are shown in Table S1.

**Table S1.** Hydrodynamic diameter (D_H_) and zeta potential (ζ) values of bare (SiO_2_) and SiO_2_-ZW nanoparticles. PDI: Polydispersity Index

| **Sample** | **D_H_ (nm)** | **PDI** | **ζ (mV)** | **D_STEM_ (nm)** |
| --- | --- | --- | --- | --- |
| **Pristine SiO_2_** | 40.7 | 0.11 | -56.6 | 30 |
| **SiO_2_-ZW** | 42.5 | 0.10 | -41.3 | 29 |

### **SECTION S3: Protein Corona Experiments**

#### *Dynamic Light Scattering (DLS)*

Figure S2 shows pristine SiO_2_ hydrodynamic diameter (D_H_) evolution with increasing BSA concentrations (in phosphate buffer, PB, at 37 °C) as determined by cumulant analysis (Z-average).

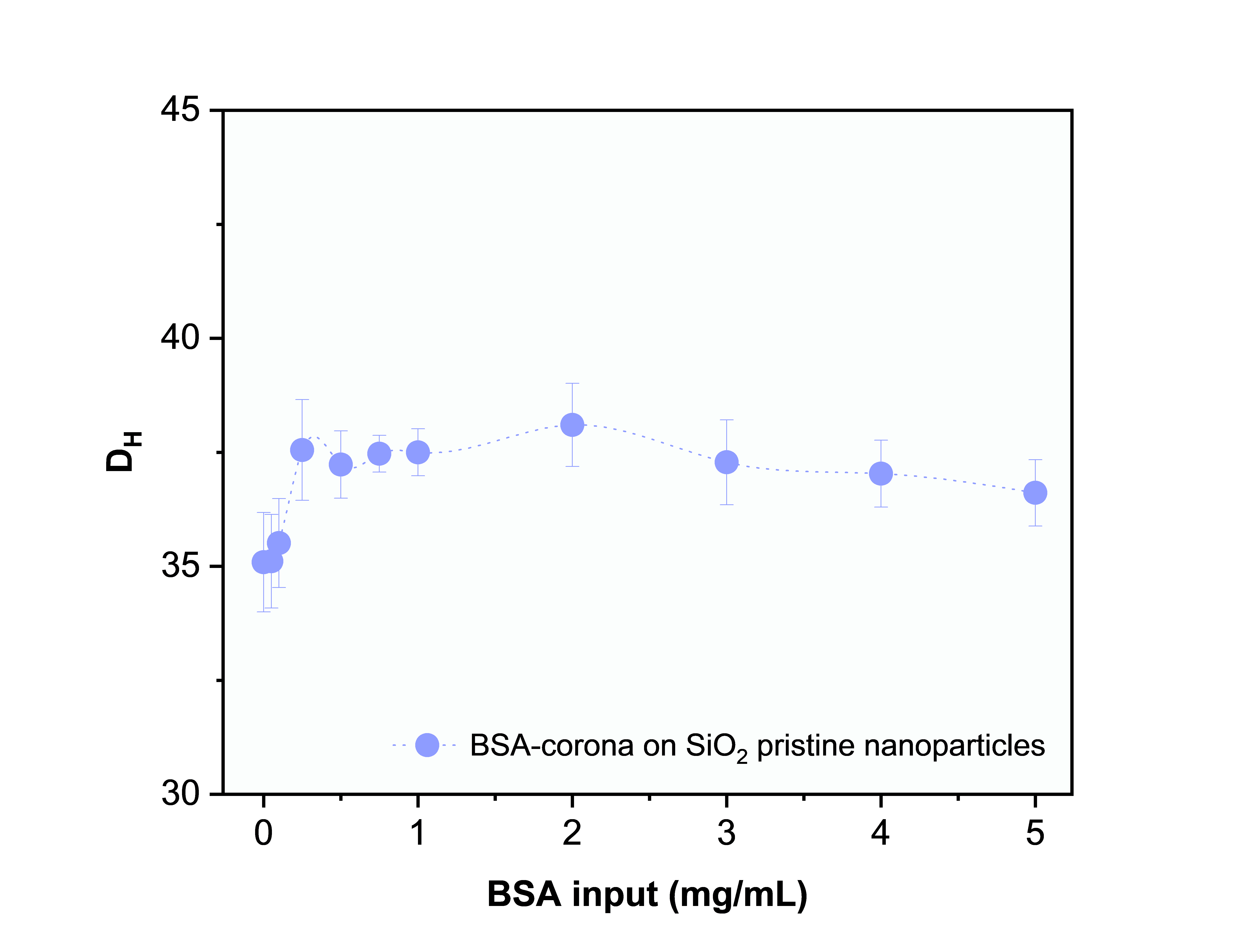

**Figure S2.** Evolution of pristine SiO_2_ (D_H_) obtained by cumulants with increasing concentrations of BSA.

#### *Synchrotron Radiation Circular Dichroism (SRCD)*

Synchrotron Radiation Circular Dichroism (SRCD) measurements were carried out at the CEDRO beamline of the Brazilian Synchrotron Light Laboratory (LNLS, Brazil) to investigate the interaction between bovine serum albumin (BSA) and SiO_2_. The experiments were performed using an OLIS DSM20 UV/VIS spectropolarimeter (OLIS, USA) operated under continuous nitrogen purge. CD spectra were recorded in the wavelength range of 178–280 nm, with a step size of 1.0 nm, an integration time of 2 s, and a constant temperature of 20 °C, using demountable quartz short-pathlength cells (0.2 mm). Three consecutive scans were collected for both the samples and the baselines. Data processing was conducted using the CDtoolX software. The analysis included averaging of scans, baseline subtraction, correction in the 265–270 nm region, and spectral smoothing using a Savitzky–Golay filter. For sample preparation, unfunctionalized silica nanoparticles were incubated with BSA at concentrations of 2 mg/mL and 4 mg/mL in PB (10 mM, pH 7.4). The incubation was performed at 37 °C for 10 min under agitation at 500 rpm in a temperature-controlled thermoblock. After incubation, the nanoparticle–protein corona (NP-PC) complexes were purified by centrifugation at 12.000 rpm and 4 °C. This purification step was repeated three times; in each cycle, the supernatant containing unbound proteins was completely discarded, and the nanoparticles were resuspended in fresh phosphate buffer. The presence of residual protein in the supernatant was monitored by UV absorbance measurements, confirming that no significant amount of unbound protein remained after the third washing step.

#### *Size-Exclusion Chromatography Coupled with Small-Angle X-ray Scattering (SEC-SAXS)*

SEC-SAXS experiments were performed at beamline B21 of the Diamond Light Source synchrotron facility (Oxfordshire, United Kingdom). Samples were loaded onto a Superdex™ 200 Increase 10/300 GL size exclusion chromatography column (Cytiva) in 20 mM HEPES pH 7.5, 150 mM KCl at 0.5 mL/min using an Agilent 1200 HPLC system. The column outlet was fed into the experimental cell, and SAXS data were recorded at 12.4 keV, detector distance 3.7 m, and each frame was exposed to the X rays radiation for 3 s. Data processing and analysis were performed using a JAVA-based program called ScÅtter, which allowed to process integrated and normalized data and perform background subtractions.

### **SECTION S4: *In Situ* Quantification of Protein Corona Through SAXS**

#### *Quantification of Adsorbed Protein in SiO_2_*

This comprehensive study incorporated two distinct protein sources: bovine serum albumin (BSA), and human serum (HS), all meticulously chosen to represent a range of biological relevance and complexity. The examination of nanoparticle-protein interactions was carried out with the utmost precision through high-throughput SAXS experiments, conducted in a controlled environment provided by the B21 beamline.^4^ Each mixture was crafted according to a standardized and pre-defined protocol, which involved the combination of equal volumes of the respective components within each individual well of a 96-well plate. The first component consisted of a fixed mass of SiO_2_ at a concentration of 4 mg/mL, suspended in water. The second component was a 2x protein solution in a 20 mM PB buffer (pH = 7.4). The combination of these two components resulted in a mixture of SiO_2_ at a concentration of 2 mg/mL, with varying protein concentrations ranging from 0 to 5 mg/mL for BSA and 0 to 5% for HS. Prior to SAXS analysis, each sample underwent a thorough homogenization process. This procedure involved subjecting the samples to 8 minutes of mixing at 450 rpm and maintaining a constant temperature of 37 °C using a ThermoMixer C (Eppendorf). This rigorous homogenization step is critical to ensure the reliability and consistency of the subsequent SAXS analysis. These measurements were performed in a beamline configuration with a beam energy of 13 keV, a sample-detector distance of 3.7 m, and a scattering vector q range of 0.0045 - 0.34 Å^-1^. The 96-well plates were arrayed in a sample environment provided by the Arinax BIOSAXS sample-handling robot,^5^ that precisely collected the liquid samples from their specific positions in the plates and injected them into the capillary (1.6 mm diameter, 10 μm thick quartz capillary held in vacuum, 50 μL of sample volume) for further analysis. In the same sample environment, PB buffer was placed in a plastic tube together with the 96-well plate to wash the capillary after each analysis to avoid accumulation of nanoparticles or proteins that could impair data acquisition. In addition, after each set of experiments the capillary was treated with Hellmanex detergent to ensure that thorough cleaning was done and that remnants of previous samples did not remain in the capillary. During the experiments, the sample environment was kept at 37 ºC, under controlled flow conditions.

#### *Step-by-step Concept for Determining Adsorbed Protein from SAXS Data*

1. Load reference SAXS datasets:

- Data1: scattering curve for bare SiO_2_ (no protein present).

- Data2: scattering curve for protein-only reference sample.

- Data3: scattering curve for the mixture (SiO_2_ + protein).

- Known total protein concentration in the mixture is denoted as pc (in mg/mL).

2. Scale the protein-only curve:

- Multiply the protein-only intensity by PC and an empirical calibration factor so that the intensity matches the scale of the mixture data.

- This scaling ensures that the protein-only curve represents the scattering from pc mg/mL of free protein in solution.

3. Interpolate curves:

- Use interpolation to generate smooth functions for the SiO_2_ and protein curves, allowing both to be compared at the same q-points.

4. Select an appropriate *q*-window:

- Choose a high-q range (e.g., 0.07 – 0.09 Å^-1^) where the scattering is dominated by free protein in solution.

- In this region, the contribution from the adsorbed corona is minimal.

5. Model the mixture scattering:

- Assume the total scattering from the mixture is the sum of the bare SiO₂ curve and a scaled protein-only curve:

I_mix_(*q*) ≈ I _SiO2_ (*q*) + s × I_protein_(*q*)

where s is the fraction of total protein that remains free in solution.

6. Fit the free-protein fraction:

- Perform a least-squares fit in the selected high-q window to determine the value of s.

- Interpretation:

s = 1.0 → all protein is free in solution (no adsorption).

s = 0.8 → 80% free, 20% adsorbed onto nanoparticles.

7. Calculate adsorbed protein concentration:

- Use the formula:

C_ads_ = (1 – s) × pc

Example: if s = 0.82 and pc = 5 mg/mL, then C_ads_ ≈ 0.90 mg/mL.

8. Present the results:

- Report the adsorbed protein concentration in mg/mL.

- Plot this value alongside the SAXS curves for visual correlation.

### **SECTION S4. Selected SAXS curves**

SAXS curves of the SiO_2_-BSA mixtures in PB, at the complete concentration range investigated (0.25-5.0 mg/mL) are presented in Figure S3.

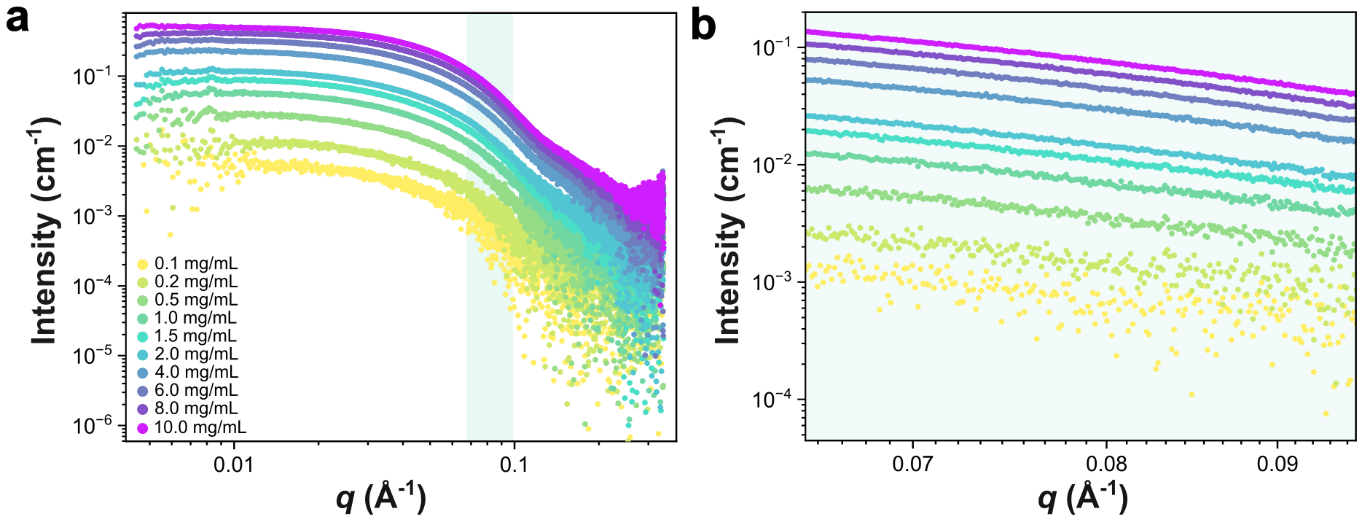

**Figure S3.** SAXS curves BSA in phosphate buffer (PB) at different BSA concentrations: 0.1, 0.2, 0.5, 1.0, 1.5, 2.0, 4.0, 6.0, 8.0 and 10.0 mg/mL. (a) Full scattering profiles over the investigated *q*-range. (b) Magnification of the high-*q* region highlighting concentration-dependent differences in scattering intensity.

Figure S4 shows the SAXS profiles of SiO_2_-human serum mixtures in phosphate buffer (PB) across the full concentration range examined (1 - 5%).

Figure S5 presents the SAXS profiles of SiO_2_-ZW mixed with BSA over the full concentration range assessed (0.25 - 5.0 mg/mL).

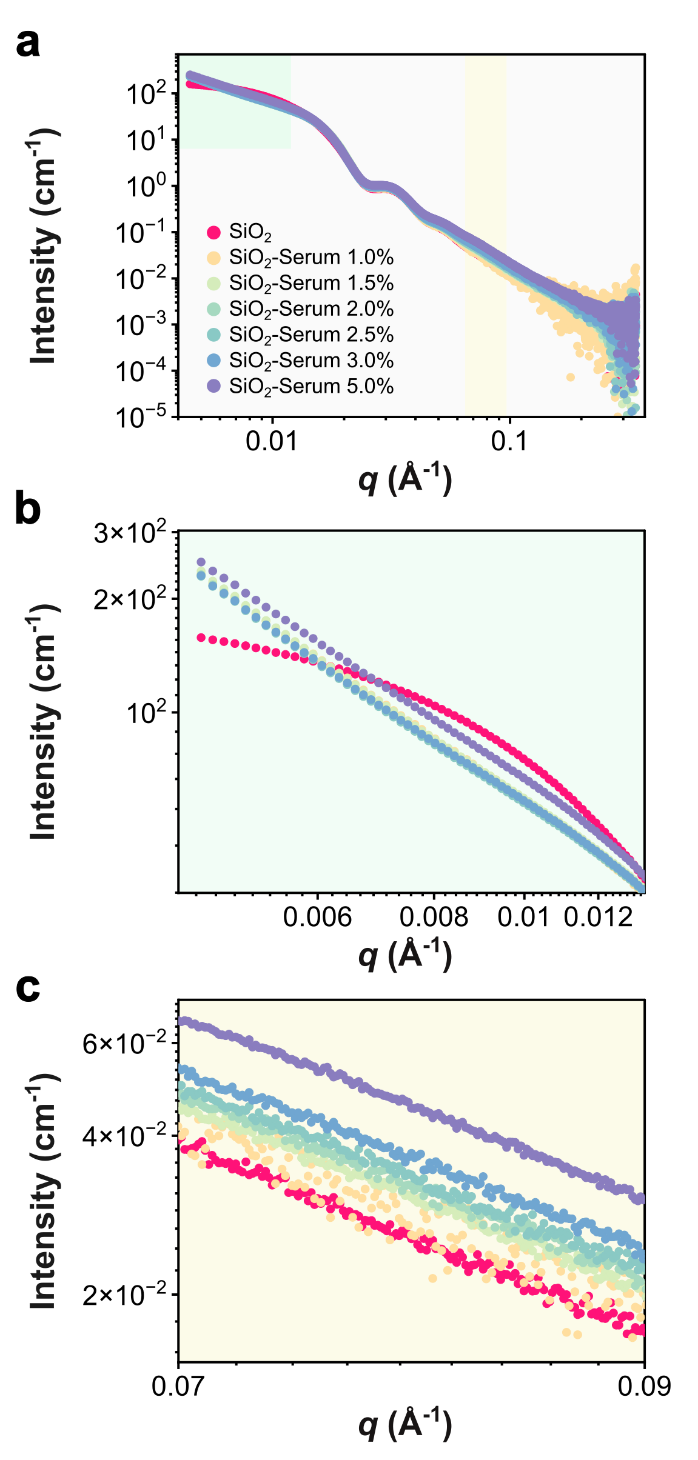

**Figure S4.** SAXS curves of mixtures of SiO_2_ nanoparticles with Human Serum (SiO_2_-Serum, respectively) in increasing protein concentration (v/v): 1.0 - 5.0 %. (a) full scattering profiles, (b) magnification of the low-*q* region highlighting aggregation profile, and (c) magnification of the high-*q* region highlighting concentration-dependent differences in scattering intensity.

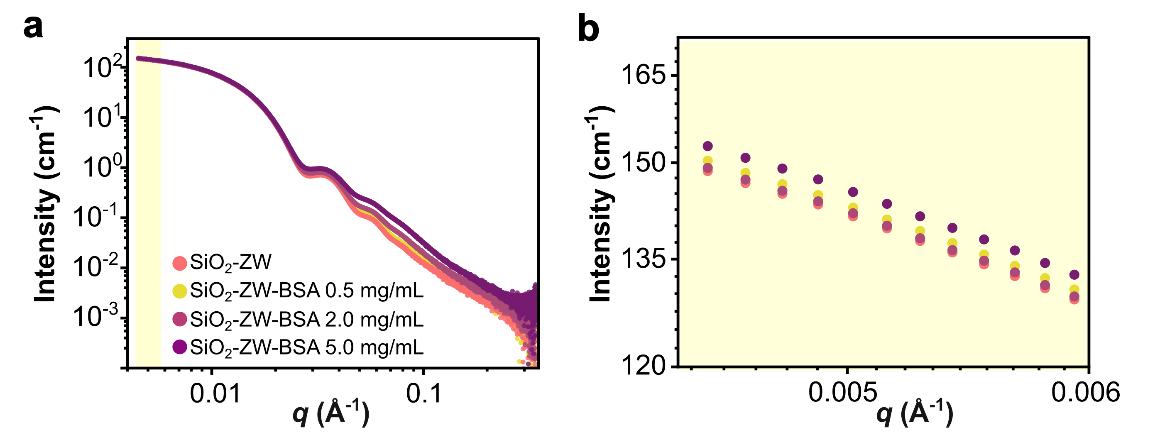

**Figure S5.** (a) SAXS profiles for SiO_2_-ZW incubated with increasing BSA concentrations (0.5, 2.0, and 5.0 mg/mL), showing minimal changes in the low-q region. (b) Magnification of the low-q region from (a), evidencing the absence of significant intensity increase even at high BSA concentrations.

### **SECTION S5. Modeling of SAXS Curves**

#### *Methodology for Generating the Heatmap of Scattering Data Variation Across q-range*

This analysis evaluates the reproducibility and variation across repeated scattering measurements by constructing a normalized deviation heatmap. The script follows these steps:

1. Data Loading: Multiple text files containing tabulated data with three columns (scattering vector (*q*), intensity (*I*), and error estimate) are loaded using a wildcard argument passed from the command line. The first three lines of each file are skipped. The q, I, and error values are extracted and stored in separate lists.
2. Normalization: For each intensity array, the total intensity (sum over all points) is computed. Then, the average total intensity across all datasets is calculated. A normalization factor for each dataset is obtained by dividing its total intensity by this average. Each dataset is then divided by its respective normalization factor, ensuring all curves are on the same relative scale.
3. Average Curve Calculation: The normalized intensity arrays are averaged pointwise to generate a mean intensity curve. This serves as a reference for subsequent variation analysis.
4. Statistical Spread (Difference) Calculation: All original (non-normalized) intensity datasets are transposed so that each column corresponds to a fixed *q* point across measurements. The pointwise minimum, maximum, and their difference are computed to quantify variability. These differences are expressed as a percentage of the mean curve and saved to a text file.
5. Visualization of Individual Curves: All raw intensity curves are plotted on a logarithmic scale, along with the average curve, to visually compare their profiles.
6. Deviation Heatmap Construction: A subset of the *q*-range is selected using index slicing (nini and nfin). For each dataset, the squared deviation from the mean curve is computed within this region. These deviations are normalized by the mean and stacked to form a 2D array.
7. Plotting the Heatmap: The deviation matrix is visualized using imshow with a color map (jet). The x-axis corresponds to *q*-points and the y-axis to different measurements. A colorbar indicates the magnitude of deviation. Custom tick labels for the x-axis are generated based on the actual *q*-values, improving interpretability.

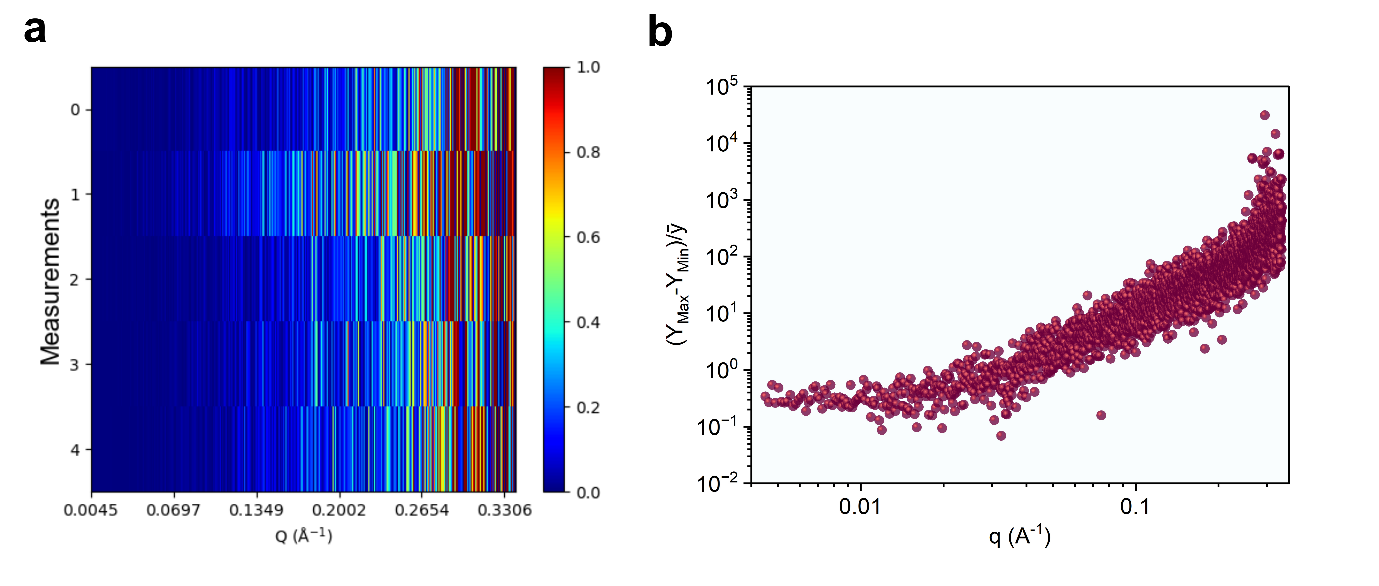

**Figure S6.** Assessment of intensity stability across the full $q$-range in SAXS measurements.
(a) Heatmap showing the variation in scattering intensity for five consecutive measurements of the same sample. (b) Plot of the relative deviation $(I_{\text{max}}-I_{\text{min}})/\bar{I}$as a function of $q$, illustrating the maximum fluctuation in intensity across the repeated acquisitions.

### **SECTION S6. Protein-nanoparticle binding isotherms**

In these experiments, the nanoparticle is systematically titrated with increasing concentrations of protein. Regardless of the specific model used, these isotherms provide valuable insights into the characteristics of the coronas formed. Specifically, we applied Langmuir, Fruendlich and Hill isotherm models to describe the experimental data.

**Table S2**: Experimental sorption capacity (Q_exp_), isotherms parameters provided by two-parameters isotherm models, with χ^2^ and coeficient of deviation (*R*^2^) error evaluation, for adsorption of BSA and HS in pristine silica nanoparticles.

|  | **BSA** | **Human Serum** |
| --- | --- | --- |
| *Q_exp_* (mg/mL) | 0.20 ± 0.02 | 0.96 ± 0.37 |
| **Langmuir** |  |  |
| Q*_ma x_* (mg/mL) | 0.23 ± 0.01 |  |
| *B* (mL/mg) | 1.61 ± 0.29 |  |
| *K_D_* (mg/mL) | 0.61 |  |
| *R_L_* | 0.13 |  |
| *R*^2^ | 0.9837 |  |
| χ^2^ | 1.2 × 10^-4^ |  |
| **Fruendlich** |  |  |
| K*f* | 0.13 ± 0.01 | 0.75 ± 0.04 |
| *n* | 2.64 | 3.01 |
| *R*^2^ | 0.9081 | 0.7649 |
| χ^2^ | 5.2 × 10^-4^ | 1.23 × 10^-2^ |
| **Hill** |  |  |
| *Q_Hill_* (mg/mL) | 0.22 ± 0.01 mg/mL | 1.59 ± 1.80 mg/mL |
| *n* | 1.23 ± 0.24 | 0.64 ± 0.66 |
| *K* (mg/mL)^n^ | 0.44 ± 0.17 | 1.05 ± 2.4 |
| *K_D_* (mg/mL) | 0.5 |  |
| *R*^2^ | 0.9862 | 0.9106 |
| χ^2^ | 1.21 × 10^-4^ | 1.17 × 10^-2^ |
